# Base editors for citrus gene editing

**DOI:** 10.1101/2021.10.19.464826

**Authors:** Xiaoen Huang, Yuanchun Wang, Nian Wang

**Affiliations:** Citrus Research and Education Center, Department of Microbiology and Cell Science, Institute of Food and Agricultural Sciences, University of Florida, 700 Experiment Station Road, Lake Alfred FL, 33850, USA

**Keywords:** Citrus, base editing, adenine base editor, cytosine base editor, *Xanthomonas*

## Abstract

Base editors, such as adenine base editors (ABE) and cytosine base editors (CBE), provide alternatives for precise genome editing without generating double-strand breaks (DSBs), thus avoiding the risk of genome instability and unpredictable outcomes caused by DNA repair. Precise gene editing mediated by base editors in citrus has not been reported. Here, we have successfully adapted the ABE to edit the TATA box in the promoter region of the canker susceptibility gene *LOB1* from TATA to CACA in grapefruit (*Citrus paradise*) and sweet orange (*Citrus sinensis*). TATA-edited plants are resistant to the canker pathogen *Xanthomonas citri* subsp. *citri* (*Xcc*). In addition, CBE was successfully used to edit the acetolactate synthase (ALS) gene in citrus. ALS-edited plants were resistant to the herbicide chlorsulfuron. Two ALS-edited plants did not show green fluorescence although the starting construct for transformation contains a GFP expression cassette. The *Cas9* gene was undetectable in the herbicide-resistant citrus plants. This indicates that the ALS edited plants are transgene-free, representing the first transgene-free gene-edited citrus using the CRISPR technology. In summary, we have successfully adapted the base editors for precise citrus gene editing. The CBE base editor has been used to generate transgene-free citrus via transient expression.

## INTRODUCTION

Unlike classical CRISPR systems that use Cas proteins, such as Cas9 and Cas12, nickase Cas9 (nCas9) derived base editors do not create double-strand breaks (DSBs). DSBs introduced by Cas proteins may pose the risk of genome instability and unpredictable outcomes caused by Non-homologous end joining (NHEJ) DNA repair mechanisms. Base editors provide alternative tools for precise genome editing without generating DSBs. Base editors are derived by tethering deoxynucleoside deaminase to a nCas9–gRNA complex that induces efficient and direct base substitutions in the genomic sequence (Rees and Liu 2018). Among the available base editors, cytosine base editors (Nishida et al. 2016; Komor et al. 2016) and adenine base editors (Gaudelli et al. 2017) enable highly efficient and precise base substitutions in a narrow window of gRNA-targeting sites. Specifically, adenine base editors mediate the conversion of A•T to G•C, whereas cytosine base editors enable the conversion of C•G to T•A in genomic DNA. It is well known that base editors can introduce specific amino acid changes in a protein, thus can be used for site-specific mutagenesis. They can also be deployed to disrupt gene functions by altering splicing sites (splice donor, splice acceptor, and branch point). CBEs can introduce premature stop codons to knock out genes. Both ABEs and CBEs can modify cis-regulatory elements to fine-tune gene functions. They can also be utilized to mutate start codon ATG to interrupt protein translation (Kluesner et al., 2021; Molla et al., 2021).

The development of base editing stemmed from seminal studies in 2016 and 2017 (Komor et al., 2016; Nishida et al., 2016; Gaudelli et al., 2017). Since then, base editing has been applied to different fields of life science including plants. Base editors have been adopted in different plant species, including *Arabidopsis* (Kang et al., 2018; Xue et al., 2018; Bastet et al., 2019; Li et al., 2019b), rice (Li et al., 2017; Lu and Zhu, 2017; Shimatani et al., 2017; Hua et al., 2018; Li et al., 2018; Xue et al., 2018; Yan et al., 2018; Zong et al., 2018; Hua et al., 2019; Li et al., 2019a; Li et al., 2020b; Ren et al., 2021), maize (Zong et al., 2017), wheat (Zong et al., 2017; Li et al., 2018; Zhang et al., 2019), tomato (Shimatani et al., 2017; Veillet et al., 2019a; Hunziker et al., 2020; Veillet et al., 2020), potato (Zong et al., 2018; Veillet et al., 2019b; Veillet et al., 2019a; Veillet et al., 2020), *Nicotiana benthamiana* (Wang et al., 2021), soybean (Cai et al., 2020), rapeseed (Kang et al., 2018; Wu et al., 2020; Cheng et al., 2021), cotton (Qin et al., 2020), watermelon (Tian et al., 2018), strawberry (Xing et al., 2020), apple (Malabarba et al., 2021), pear (Malabarba et al., 2021), and poplar tree (Li et al., 2021a). Base editors have not been reported in citrus. Previously, CRISPR/Cas has been successfully used in genome editing of citrus (Jia et al., 2017; Zhang et al., 2017; Jia et al., 2019b; Zhu et al., 2019; Huang et al., 2020; Jia and Wang, 2020; Huang et al., 2022) despite the challenges in citrus transformation owing to its recalcitrant nature. Importantly, we have developed a very efficient, improved CRISPR/Cas9 system for citrus genome editing (Huang et al., 2022), which we took advantage of for the base editors in citrus in this study.

Citrus is one of the most important fruit crops in the world and faces many disease challenges including citrus Huanglongbing and citrus canker (Gochez et al., 2018; Wang, 2019). Citrus canker is caused by *Xanthomonas citri* subsp. citri (*Xcc*). Most commercial citrus varieties, including grapefruit and sweet orange varieties, are susceptible to canker disease. *Xcc* causes the characteristic hypertrophy and hyperplasia symptoms on citrus tissues via secretion of PthA4, a transcriptional activator-like (TAL) effector, through the type III secretion system (Swarup et al., 1992; Hu et al., 2014). PthA4 enters the nucleus and activates the expression of the canker susceptibility (S) gene *LATERAL ORGAN BOUNDARIES 1* (*LOB1*) via binding to the effector binding elements (EBE) in the promoter region (Hu et al., 2014). In previous studies, canker-resistant citrus plants were generated by editing the EBE or the coding region of *LOB1* (Jia et al., 2019b; Jia and Wang, 2020; Jia et al. 2022; Huang et al., 2022). Intriguingly, in most cases, *Xanthomonas* TAL EBE in the promoter of S genes overlaps with or locates immediately downstream of the TATA box of the S genes (Huang et al., 2017). TATA box is a core promoter element conserved both in plants and animals, with the consensus sequence TATA(A/T)A(A/T). The TATA box is pivotal in transcriptional activation. The TATA box of the *CsLOB1* promoter is overlapped with the EBE region. In this study, we aimed to test if the TATA box can be edited with ABE8e (Richter et al., 2020b). We reasoned that editing of the EBE-associated TATA box may abolish or reduce the induction of S genes by *Xanthomonas* TAL effectors to generate *Xanthomonas*-resistant crops.

Unlike most economically important crops, citrus species reproduce through apomixis. Apomixis is a way of asexual reproduction with offspring genetically identical to the mother plant (Wang et al., 2017b). Apomixis facilitates fixing desired traits, hybrid vigor and heterozygosity. However, one of the disadvantages of apomixis is the lack of sexual crosses, hence the lack of genetic segregation in the next generation of citrus. Therefore, it is challenging to obtain transgene-free, gene-edited citrus through genetic segregation. Transgene-free gene-edited crops such as rice, maize, wheat, are usually obtained through genetic segregation in the next generation. In addition, fruit trees such as citrus, have a long juvenile period (5-10 years). Thus, it is crucial to generate transgene-free gene-edited citrus in the T0 generation. In this study, we explored the possibility to obtain transgene-free, gene-edited citrus through base editors, such as CBE.

## MATERIAL AND METHODS

### Making ABE construct

The binary vector backbone PC-35S was first modified to contain a CsU6-tRNA-gRNA scaffold cassette (Huang et al., 2020) with two *Aar*I sites for the gRNA insertion. The vector also contains a unique *Xba*I site downstream of the 35S promoter and a unique *Eco*RI site right upstream of the HSP terminator. Dicot plant codon-optimized Cas9 gene from the pXH1 vector (Huang et al., 2020) was first mutated at D10 to A to make Cas9^D10A^ nickase or nCas9. The evolved TadA8e (Richter et al., 2020b) was PCR amplified using the ABE8e plasmid (Addgene) as template and primers ABE8-F1/R1; nCas9 was PCR amplified using primers ABE8-F2/R2 and pXH1 as template. The modified PC-35S was digested with *Eco*RI+*Xba*I Ligation of TadA8e, nCas9, and *Eco*RI/*Xba*I-digested vector was performed using the in-fusion cloning method (Takara Bio) to make vector PC-ABE8e. The CmYLCV promoter (Huang et al., 2022), which confers high gene editing efficiency in citrus, was PCR amplified with primers CmY-F2/CmY-R2. The 35S promoter in PC-ABE8e was replaced with the CmYLCV promoter to make the final vector PC-CmYLCV-ABE8e. Primers LOBBE-F1/ LOBBE-R1 (for gRNA GTTTATATAGAGAAAGGAAA) were annealed and cloned into *Aar*I-digested PC-CmYLCV-ABE8e. All constructs were verified with Sanger sequencing.

### Making CBE construct

The PC-ABE8e vector described above was digested with *Sbf*I+*BspE*I to remove TadA8e. The CmYLCV promoter (Huang et al., 2022) was PCR amplified using primers CmY-F1/R3. The fragment A3A-RAD51DBD (Supporting information 1) was synthesized by Integrated DNA Technologies, Inc. (Coralville, IA, USA). Ligation of the CmYLCV promoter, A3A-RAD51DBD, and *Sbf*I/*Bsp*EI-digested vector was performed using the in-fusion cloning method (Takara Bio) to make an intermediate vector CmYLCV-A3A-RAD51-nCas9. CmYLCV-A3A-RAD51-nCas9 was further digested with *Eco*RI. The UGI was PCR amplified using primers UGI-F1/R1 and cloned into *Eco*RI site of the CmYLCV-A3A-RAD51-nCas9 vector via the in-fusion cloning method (Takara Bio) to construct the final CBE vector. Two gRNAs for two different alleles of the citrus *ALS* gene were designed. Primers ALS-F/ALS-R were used to amply gRNA scaffold-tRNA unit using the plasmid pXH1 (Huang et al., 2020) as template. The amplicon was digested with *Bsa*I and cloned into the *Aar*I-digested CBE to make the CBE-2xALS construct. All constructs were verified by Sanger sequencing.

### Citrus transformation

The constructs were transformed into *Agrobacterium* strain EHA105. *Agrobacterium*-mediated transformation of citrus epicotyl was performed as described previously (Jia et al., 2019a; Huang et al., 2020; Huang et al., 2022). For the selection of herbicide chlorsulfuron resistant citrus, citrus epicotyl segments were cultured on kanamycin-containing selection media for one week. After one week, the citrus epicotyl segments were transferred to chlorsulfuron (Fisher Scientific, Catalog No.50-255-082) containing media (150 nM) without kanamycin. Every three weeks, the citrus epicotyl segments were transferred to new chlorsulfuron-containing media to select chlorsulfuron-resistant shoots.

### Genotyping of citrus transformants

We performed genotyping of citrus transformants as described previously (Huang et al., 2022). Citrus genomic DNA was extracted using the CTAB (cetyltrimethylammonium bromide) method. Detection of editing in the target genes was performed via amplifying the target regions (primers in Supplemental Table 1) with high fidelity DNA polymerase Q5 (New England Biolabs, Ipswich, MA, USA), followed by cloning of PCR products and sequencing. Primers LOBpro-F1/ LOBpro-R1 were used for the *LOB1* promoter genotyping. Primers CsALSgt-F2/ CsALSgt-R2 were used for *ALS* genotyping. Primers Cas9gt-F1/ Cas9gt-R1 were used for *Cas9* genotyping. Primers CsALSgt-F1/CsALSgt-R1 were used for the PCR detection of *ALS*.

### *Xcc* inoculation

*Xanthomonas citri* subsp. *citri* (*Xcc*) wild type strain 306 and dLOB2 containing *Xcc pthA4*:Tn5 (Hu et al., 2014; Teper et al., 2020; Huang et al., 2022) were suspended in 20 mM MgCl2 at 10^7^ CFU/ml. The bacterial suspensions were syringe-infiltrated into fully expanded young leaves of wild type grapefruit, sweet orange Hamlin plants, or edited lines. Inoculated plants were kept in a temperature-controlled (28°C) glasshouse with high humidity. Pictures were taken eight days post inoculation for disease resistance evaluation.

### Analysis of potential off-targets

To analyze potential off-targets, we analyzed the putative off-targets using a web-based software (http://crispr.hzau.edu.cn/cgi-bin/CRISPR2/CRISPR). Genomic DNA from transgenic lines was used as template, and the primers listed in Table S1 were used to amplify the fragments spanning the off-targets. Finally, the PCR products were subjected to Sanger sequencing.

## RESULTS

### Citrus optimized ABE8e construct can precisely edit citrus genes in transient assays

To test if precise gene editing works in citrus, we first tested adenine base editors (ABE), ABE8e which mediates A•T-to-G•C base changes (Richter et al., 2020b). The citrus optimized ABE8e vector was constructed (Figure 1A). TadA8e (Richter et al., 2020b) was N-terminally fused with Cas9^D10A^ nickase. There is a nuclear localization signal (NLS) at each end of TadA8e-Cas9^D10A^ to increase its nucleus transportation. The CsU6-tRNA-gRNA-scaffold unit for multiplex editing in this ABE system was described previously (Huang et al., 2020). We sought to edit the TATA box (Figure 1B, D) located upstream of the EBE of the promoter of the citrus canker susceptibility (S) gene *LOB1* with ABE8e. A nearby NGG PAM site enables the TATA box within the editing window of ABE8e. The EBE region of the *LOB1* promoter in citrus is responsible for binding by the TAL effector PthA4 of *Xcc* (Hu et al., 2014) to activate its expression. The TATA box of the *LOB1* promoter overlaps with the EBE region (Figure 1B, D). Hu *et. al* previously showed that mutation of TATA box abolishes *LOB1* induction by PthA4 in the transient assay (Hu et al., 2014). We first investigated if the ABE construct targeting the TATA box can edit the target as expected through *Xcc*-facilitated agroinfiltration of citrus leaves (Jia and Wang, 2014). The transient assay (Figure 1C) showed that the construct edited the TATA box as anticipated (Figure 1D, E). The ABE mutated <u>T</u>A<u>T</u>A to <u>C</u>A<u>C</u>A (from T<u>A</u>T<u>A</u> to T<u>G</u>T<u>G</u> for the complementary strand). Test on another gene *CsTub* (*Cs1g21050*) through transient assay also demonstrated precise editing (Figure 1F, G).

**FIGURE 1.**
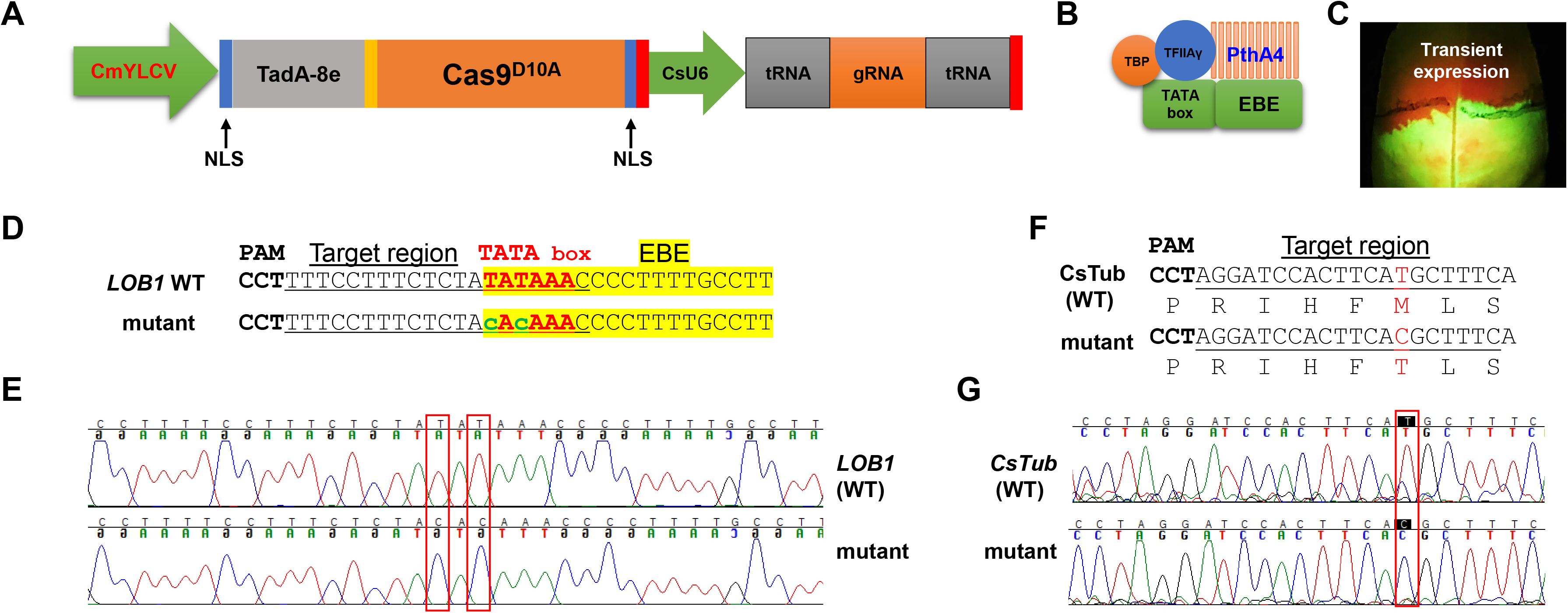
Precise gene editing in citrus with adenine base editor ABE8e via transient expression. (A) Illustration of ABE8e adenine base editing (ABE) system for citrus. CmYLCV (Huang et al., 2022), *Cestrum yellow leaf curling virus* promoter; TadA-8e (Richter et al., 2020b), evolved *Escherichia coli* tRNA adenosine deaminase; Cas9^D10A^, Cas9 nickase; NLS, nuclear localization signal; CsU6, citrus U6 promoter. (B) TATA box located upstream of the *LOB1* EBE region is associated with general transcription factors and the TAL effector PthA4 of *Xanthomonas citri* subsp. *citri* (*Xcc*). (C) *Xcc*-facilitated transient expression of the PC-CmYLCV-ABE8e-LOB1 construct in citrus leaf. The construct carries a GFP expression cassette. (D) Editing of TATA in the TATA box of the *LOB1* promoter into CACA in transient assay. Underlined nucleotides were selected for gRNA design (gRNA: GTTTATATAGAGAAAGGAAA); EBE, *Xanthomonas* TAL effector binding element, highlighted with yellow; TATA box, in red font; edited sequences, in lower case with green font. (E) Chromatograms for D), *LOB1* WT (upper), and mutant (lower). Mutation sites are indicated within red rectangles. (F) Editing of the *CsTub* gene (*Cs1g21050*) in the transient expression assay. The amino acids are aligned under the corresponding DNA sequences. The mutation of T to C changes the corresponding amino acid from methionine (M) to threonine (T). (G) Chromatograms for (E), *CsTub* WT (upper) and mutant (lower). Mutation sites are indicated within red rectangles.

### Editing the TATA box of the *LOB1* promoter confers citrus resistance to *Xcc*

Biallelic editing of *CsLOB1* coding region results in *Xcc* resistance while maintaining normal plant development and growth (data not shown). We expected that editing of the TATA box of *CsLOB1* would not affect plant development and growth either. *Agrobacterium*-mediated stable transformation of grapefruit (*Citrus paradise*) and sweet orange (*Citrus sinensis*) Hamlin epicotyl tissues was conducted to edit the TATA box in the *CsLOB1* promoter of both varieties. Two transgenic grapefruit plants and one transgenic sweet orange were obtained. Genotyping showed that in the grapefruit line #2 and Hamlin sweet orange mutant the TATA box was successfully edited (Figure 2A, B). The <u>T</u>A<u>T</u>A box in the *LOB1* promoter in grapefruit line #2 and Hamlin swet orange mutant was 100% edited into <u>C</u>A<u>C</u>A. The purity of editing was confirmed through direct sequencing of PCR products and colony sequencing of cloned PCR products. In grapefruit line #1, the first T in TATA box was 100% mutated to C, while 56% of the second T in the TATA box was mutated into C (9 clones out of 16). Another T to C editing was observed immediately upstream of TATA box in the grapefruit line #1 (31.2%, 5 clones out of 16). Using the ABE system, we achieved high rate (66.7%) of biallelic/homozygous editing of the TATA box in the *LOB1* promoter. After micro-grafting, one transgenic grapefruit plant and one transgenic sweet orange plant survived. Inoculation of the TATA-edited plants with *Xcc* demonstrated that the TATA-edited plants were resistant to *Xcc* (Figure 2C). On the contrary, all plants were susceptible to *Xcc pthA4*:Tn5 dLOB2, which carries a designer TAL dLOB2 for LOB2 induction to cause canker disease in citrus plants. *Xcc pthA4*:Tn5 dLOB2 was included in the inoculation assay as a control. The onset of canker disease by *Xcc pthA4*:Tn5 dLOB2 on the same leaves excluded the possibility that lack of canker disease on the TATA-edited mutant plants is due to leaf age. These results demonstrated that base editor ABE8e can efficiently and precisely edit the citrus genome (both grapefruit and sweet orange). These results also showed a promising strategy by editing the EBE-associated TATA box of S genes to breed *Xanthomonas*-resistant crops through base editor ABE8e.

**FIGURE 2.**
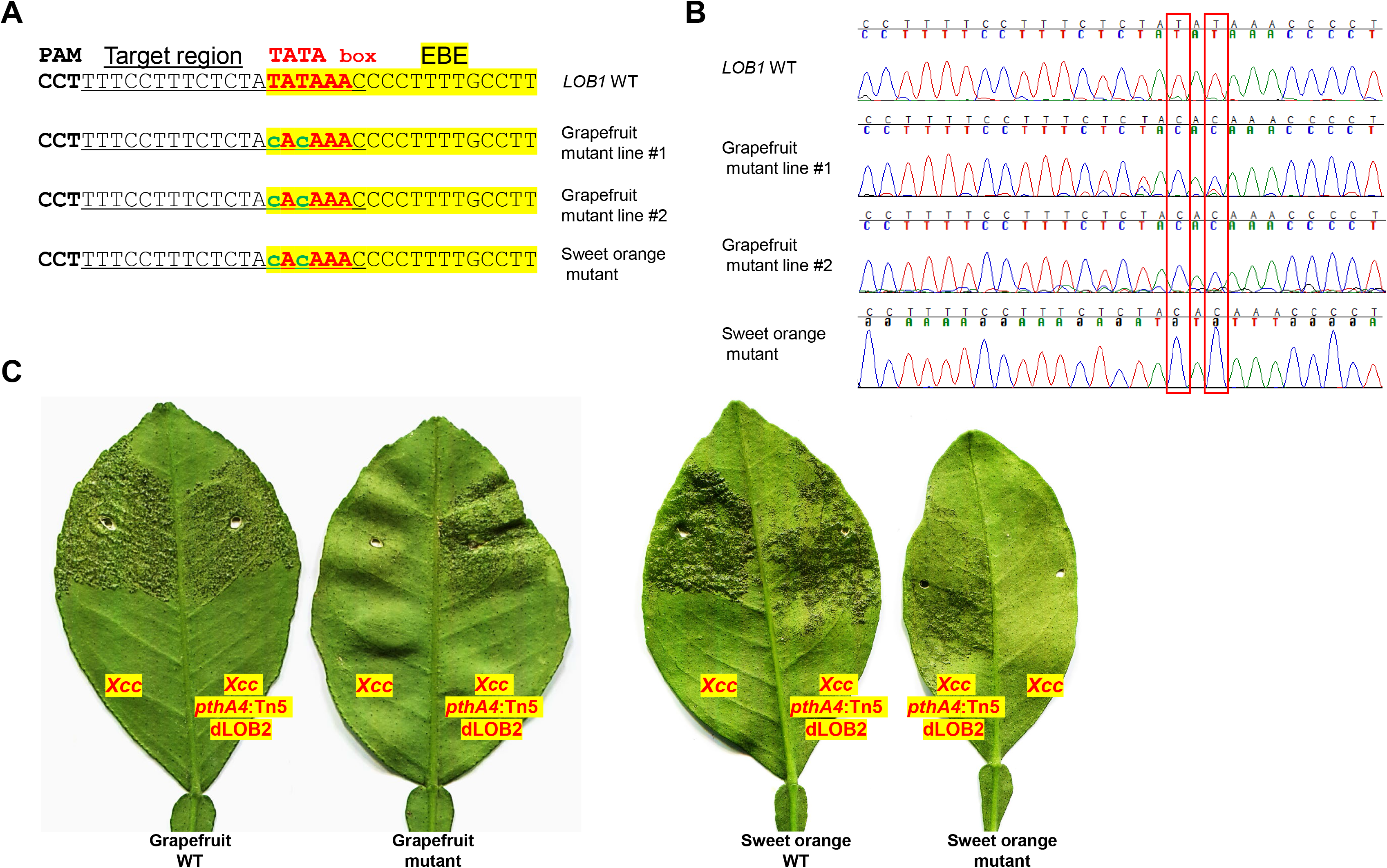
TATA-edited grapefruit (*Citrus paradise*) and sweet orange (*Citrus sinensis*) are resistant to the canker pathogen *Xanthomonas citri* subsp. *citri* (*Xcc*). (A) Edition of the TATA box into CACA in stable transgenic grapefruit and sweet orange. Underlined nucleotides were selected for gRNA design (gRNA: GTTTATATAGAGAAAGGAAA); EBE, *Xanthomonas* TAL effector PthA4 binding element, highlighted with yellow; TATA box, in red font; edited sequences, in lower case with green font. (B) Chromatograms for (A). Mutation sites are indicated within red rectangles. (C) Inoculation of WT and TATA-edited mutant plants with *Xcc* or *Xcc pthA4*:Tn5 dLOB2 in the indicated areas of leaves. *Xcc pthA4*:Tn5 dLOB2, an *Xcc pthA4* mutant strain carrying a designer TAL effector dLOB2 for the induction of citrus *LOB2* expression to cause canker symptoms. *Xcc pthA4:Tn5* dLOB2 was used as control.

### Developing a CBE for citrus

Next, we tested if cytosine base editors (CBE), which can mediate C•G-to-T•A base changes, work in citrus. A citrus optimized CBE vector was constructed (Figure 3A). We chose APOBEC3A (A3A) deaminase for citrus CBE considering its wide deamination window and high editing efficiency (Zong et al., 2018; Zhang et al., 2020). An inhibitor of uracil DNA glycosylase (UGI) was fused immediately downstream of nCas9 to increase efficacy of C to T editing. A single-stranded DNA-binding domain from Rad51 protein (RAD51-DBD) was inserted between the deaminases and nCas9, which was reported to substantially increase activity and an expand editing window of CBE (Zhang et al., 2020). Tan et al. recently reported that RAD51-DBD can also dramatically increase editing efficiency of ABE (Tan et al., 2022). This CBE system contains a CsU6-tRNA-gRNA-scaffold unit for multiplex editing. The final CBE construct also carries a GFP expression cassette for direct visualization of transgene integration. We first determined whether the citrus *acetolactate synthase* (*ALS*) gene can be edited with this CBE. Most citrus varieties, owing to their hybrid nature, have two different alleles of the *ALS* gene responsible for herbicide resistance. We designed two gRNAs targeting both *ALS* alleles. The transient expression result showed that the CBE construct edited both alleles of the *ALS* gene as expected (Figure 3B, C).

**FIGURE 3.**
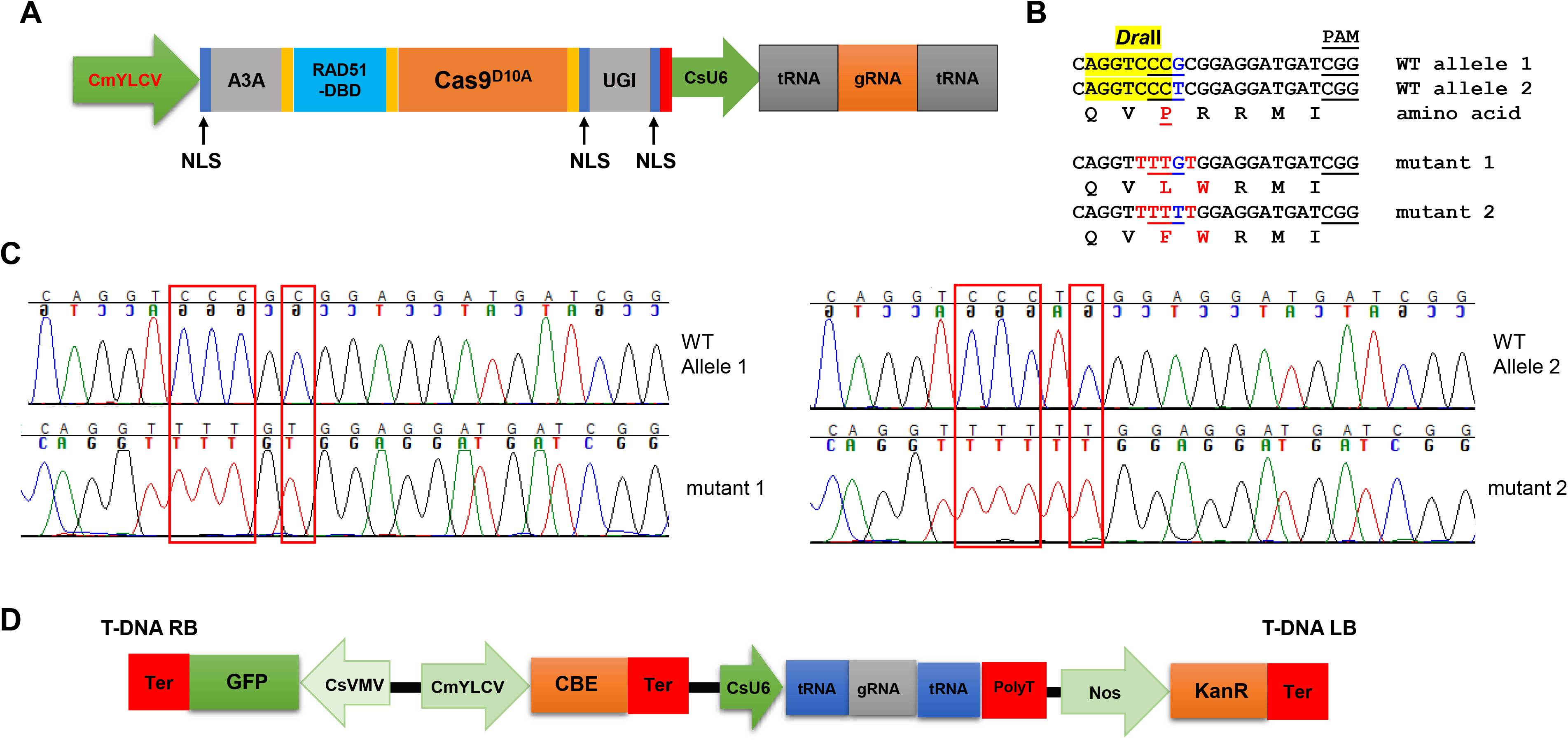
Precise gene editing in citrus with cytosine base editor A3A-RAD51-DBD via transient expression. (A) Illustration of cytosine base editing system (CBE). A3A, human APOBEC3A cytidine deaminase; RAD51-DBD, RAD51 DNA-binding domain; UGI, uracil glycosylase inhibitor; Cas9^D10A^, Cas9 nickase; NLS, nuclear localization signal; CsU6, citrus U6 promoter. (B) CBE base editing of the *ALS* gene via transient expression assay. There are two ALS alleles in citrus. Two gRNAs (gRNA1: CAGGTC<u>CCG</u>CGGAGGATGAT and gRNA2: CAGGTC<u>CCT</u>CGGAGGATGAT) were designed to edit both ALS alleles using the multiplex CBE construct. The amino acids are aligned under the corresponding DNA sequences. The restriction enzyme *Dra*II-resistant PCR amplicon was subject to cloning and sequencing. The restriction enzyme *Dra*II recognition site, highlighted in yellow. Edited sites, in red font. (C) Chromatograms for (B). Mutation sites are indicated within red rectangles.

### Developing transgene-free, herbicide-resistant citrus

Acetolactate synthase (ALS) is an enzyme required for the biosynthesis of multiple branched-chain amino acids, such as valine, leucine, and isoleucine. Chlorsulfuron is a known ALS inhibitor that kills plants, thus being used as a herbicide. A single amino acid mutation in *ALS* genes in various plants confers resistance to chlorsulfuron (Kuang et al., 2020; Malabarba et al., 2021). For example, when the amino acid P in the “QVPRRMI” amino stretch of ALS protein is mutated to S or F (Figure 3B, Figure 4C), the plant becomes resistant to chlorsulfuron. We first tested the sensitivity of citrus to chlorsulfuron. The growth of citrus Carrizo citrange, a hybrid of *Citrus sinensis* ‘Washington’ sweet orange X *Poncirus trifoliata*, was completely inhibited by chlorsulfuron (Figure 4A). *Agrobacterium*-mediated stable transformation of Carrizo epicotyl tissues was performed with chlorsulfuron selection in the culture media after one week of culture. Two chlorsulfuron-resistant plants grew on the chlorsulfuron-containing medium while most of the transformed epicotyl tissues did not grow (Figure 4B). Genotyping of the chlorsulfuron-resistant plants showed that both alleles of the *ALS* gene were edited in the two edited plants (Figure 4C, D). Only C-to-T substitutions, but no other by-product substitutions, were observed in the edited plants. The editing rendered the “QVPRRMI” amino stretch of ALS protein to “QVSRRMI” in one allele, and to “QVFWRMI” in another allele. Both mutant plants have same genotypes in *ALS* locus. Intriguingly, these two plants did not show green fluorescence although the construct used for transformation contains a GFP expression cassette (Figure 3D). To further confirm the absence of T-DNA integration, we performed PCR amplification of the *Cas9* gene in the mutant plants and the *Cas9* gene was undetectable in the herbicide-resistant citrus plants (Figure 4E). This indicates that the *ALS*-edited plants are transgene-free, which likely resulted from the transient expression of the CBE construct, representing the first transgene-free gene-edited citrus using the CRISPR technology.

**FIGURE 4.**
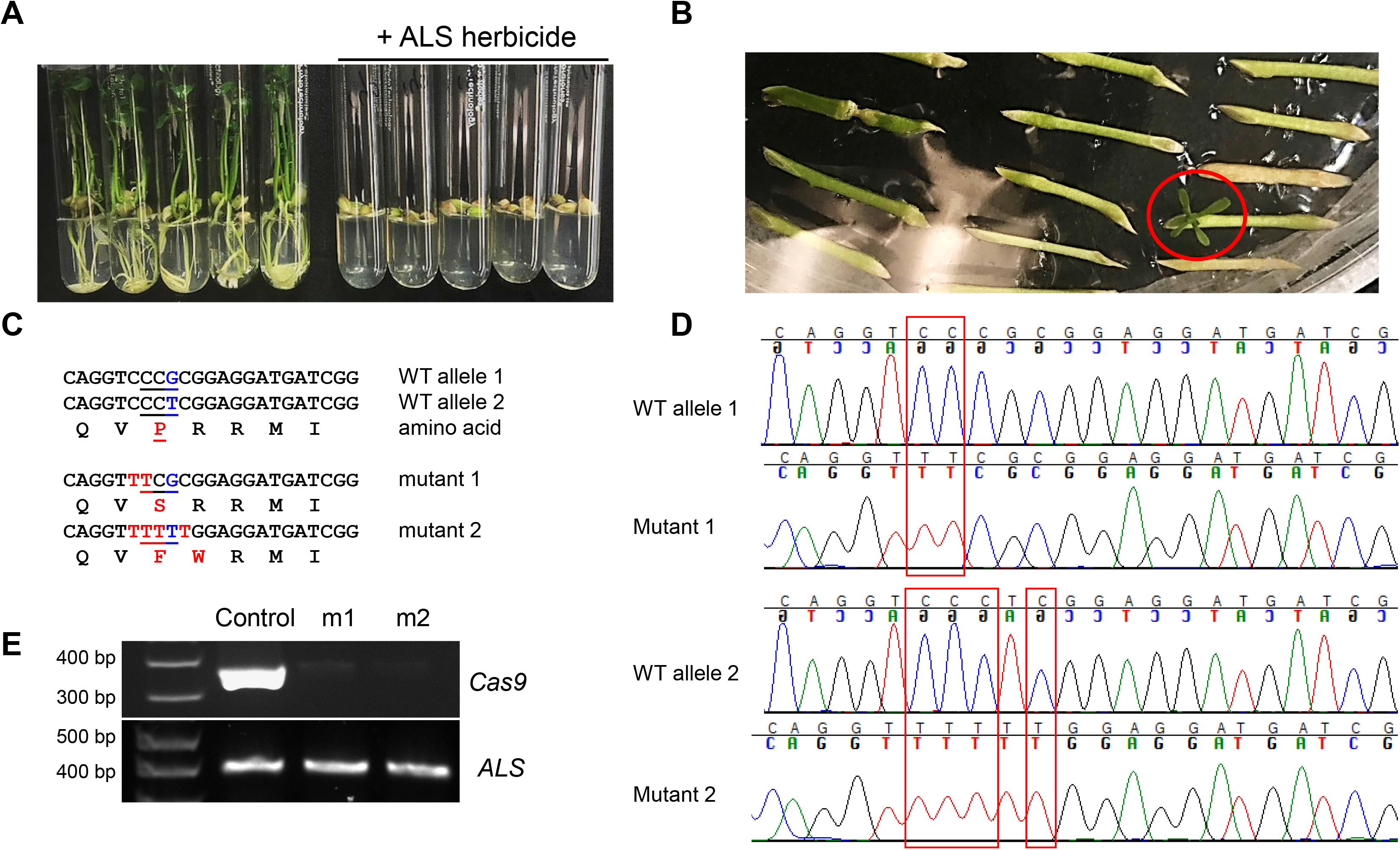
Transgene-free editing of the *ALS* gene in citrus confers herbicide resistance. (A) The growth of citrus seedlings (Carrizo citrange) was inhibited by the herbicide chlorsulfuron. (B) Selection of herbicide-resistant Carrizo citrange on chlorsulfuron-containing media (150 nM). The chlorsulfuron-resistant regenerated plant was indicated by a red circle. (C) Sequencing results from chlorsulfuron-resistant mutant plants. The amino acids are aligned under the corresponding DNA sequences. Edited sites, in red font. (D) Chromatograms for (C). Mutation sites are indicated within red rectangles. Sanger sequencing results of the PCR amplicons that were cloned for colony sequencing. For each mutant plant, 14 clones were subjected to Sanger sequencing. (E) PCR of *Cas9* and *ALS* for control plant (GFP positive) and herbicide chlorsulfuron-resistant citrus plants.

### Off-target analysis in the mutant lines

To investigate whether base editors can introduce off-target mutations, we amplified and sequenced top 12 potential off-target sites in both Grapefruit and sweet orange Hamlin TATA-edited mutant lines. The top potential off-targets all carry 4 or more mismatches. Sequencing results showed that no edits were detected at potential off-target sites (Supplemental Table 2). For *ALS*-edited plants, only 1 potential off-target exists with four or fewer mismatches within the protospacers. Sequencing results showed that no edits were detected at the potential off-target site.

## DISCUSSION

TATA box plays an important role in recruiting the basal transcription factors for assembly into transcription machinery. The TATA box is a key determinant of promoter strength (Jores et al., 2021). Previously, Hu et. al showed that mutation of TATA box in the *LOB1* promoter abolished the *LOB1* induction by TAL effector PthA4 in transient assay (Hu et al., 2014). In this study, we successfully used an ABE (ABE8e variant (Richter et al., 2020b)) to edit the TATA box of *LOB1* in citrus. Mutation of TATA in the TATA box to CACA confers resistance to citrus canker disease caused by *Xcc*. To our knowledge, this is the first report that editing the TATA box in the promoter with an ABE can confer disease resistance in plants. The TATA box in the promoter region of overexpressed PMP22 was targeted by CRISPR/Cas9 to knock down its expression level to cure the disease Charcot-Marie-Tooth 1A (CMT1A) in mice model (Lee et al., 2020). *Xanthomonas* species can infect a wide range of crops, such as rice, wheat, citrus, tomato, pepper, cabbage, cassava, banana, mango, sugarcane, cotton, bean, strawberry, and lettuce. TAL effectors secreted by *Xanthomonas* species via type III secretion system can induce S genes to cause diseases. By editing the TATA box in the S genes with an ABE, our strategy may be applied to other crops for breeding of *Xanthomonas*-resistant crops.

ABE has been artificially evolved from ABE7.10 to ABE8e, which dramatically increases deamination activity (Richter et al., 2020a; Richter et al., 2020b). ABE8e has been adapted for base editing in plants, such as in rice (Daqi Yan 1 2021; Li et al., 2021b; Ren et al., 2021; Wei et al., 2021; Xu et al., 2021). The editing efficiency varied from 33% to more than 93%. A recent report demonstrated that ABE8e achieved 30%-60% editing efficiency in *Nicotiana benthamiana* (Wang et al., 2021). In our current study, we adapted ABE8e to edit the TATA box of *LOB1* promoter in citrus and achieved 66.7% editing efficiency. We speculate that the highly efficient base editing in citrus can be attributed to both the highly efficient ABE8e and highly efficient improved CRISPR/Cas9 system that we developed (Huang et al., 2022). However, owing to the recalcitrant nature of citrus to genetic transformation, gene editing of citrus remains challenging despite the high efficacy of CRISPR/Cas9-mediated gene editing.

For many economically important crops, such as rice and maize, it is easy to obtain transgene-free gene-edited crops by simply choosing the segregating progenies that do not contain the CRISPR construct. For fruit trees such as citrus, it has been extremely difficult to obtain transgene-free gene-edited plants, which is because of the long juvenile period and reproduction of citrus through apomixis (Wang et al., 2017b). Due to the apomixis reproduction nature, citrus lacks genetic segregation in the next generation. Therefore, the CRISPR constructs in transgenic citrus cannot be segregated out like rice and maize. In this study, we successfully generated gene-edited, transgene-free citrus with a CBE. We obtained transgene-free, *ALS*-edited citrus by the selection of citrus plants on herbicide chlorsulfuron-containing media. The selection pressure exerted by herbicide chlorsulfuron facilitated the selection of *ALS*-edited plants, among which some are transgene-free through transient expression of CBE construct. A similar strategy has been reported previously in other crops (Veillet et al., 2019a).

In our current study, we used A3A-based CBE (Zong et al., 2018) to edit the citrus *ALS* gene to confer herbicide resistance. It was reported that A3A-based CBE outperformed rAPOBEC1-BE3, hAID-BE3, and PmCDA1-BE3 in base editing efficiency (Zong et al., 2018; Cheng et al., 2021; Li et al., 2021a; Randall et al., 2021). Off-target mutations caused by CBEs have been reported in tomato, rice, and mouse (Shimatani et al., 2017; Jin et al., 2019; Zuo et al., 2019; Randall et al., 2021). Although we detected no off-target mutations at predicted off-target sites, we cannot rule out that there are gRNA-independent off-target mutations. Randall et. al reported low level of sgRNA-independent off-target mutations in tomato mediated by A3A-based CBE, though the difference is not statistically significant (Randall et al., 2021). They proposed that reduced expression level and/or duration of CBE can further reduce off-target mutations. Transient expression of a CBE in tomato to edit SlALS1 greatly reduced the risk of sgRNA-dependent off-target editing at the SlALS2 locus, compared to the constitutive expression of the CBE through stable transformation (Veillet et al., 2019a).In our study, we achieved *ALS*-edited citrus with CBE through transient expression. Therefore, according to the study by Randall et. al (Randall et al., 2021), the *ALS*-edited citrus might have minimum off-target mutations. The strategy we developed in this study can transiently express CBE to achieve transgene-free genome editing and may also minimize off-target mutations. In contrast to CBE, ABEs produce much cleaner edits and do not generate genome-wide gRNA independent off-target mutations (Jin et al., 2019; Zuo et al., 2019). Noticeably, the rate of unwanted SNPs caused by ABEs is comparable to spontaneous mutations (Jin et al., 2019). Consistently, in our study we detected no off-target mutations.

It is probable to generate non-transgenic citrus varieties by simultaneously editing ALS and genes of interest by using our citrus optimized multiplex CBE and selecting the regenerated plants on herbicide chlorsulfuron-containing media. In this way, we can select transgene-free citrus with desired agronomic traits. For example, we may take advantage of ABE-CBE dual-editor (Li et al., 2020a) to simultaneously edit *CsLOB1* EBE and *CsALS* to select transgene-free, canker-resistant citrus. It is challenging for many vegetatively propagated crops and hybrid crops to obtain transgene-free gene-edited plants. The strategy that we discussed here may also be applicable to other vegetatively propagated crops and hybrid crops to obtain transgene-free plants with desired traits.

In summary, we have successfully adapted base editors for citrus gene editing and have generated transgene-free gene-edited citrus plants. Such tools will be useful to tackle the challenges the citrus industry is facing, such as Huanglongbing (HLB) (Wang et al., 2017a; Wang, 2019).

## Abbreviations

ABE: adenine base editor
CBE: cytosine base editor
CRISPR: clustered regularly interspaced short palindromic repeats
EBE: TAL effector-binding element
gRNA: guide RNA
*Xcc*: *Xanthomonas citri* subsp. *citri*

## Acknowledgments

The research has been supported by USDA National Institute of Food and Agriculture grant # 2018-70016-27412, #2016-70016-24833, and #2019-70016-29796, USDA-NIFA Plant Biotic Interactions Program 2017-67013-26527, Florida Citrus Initiative, and Florida Citrus Research and Development Foundation.

## Conflicts of interest

The authors declare no competing financial interests.

## Author contributions

XH designed and performed the experiments. YW performed micro-grafting and off-target experiments. XH and NW wrote the paper. NW supervised the study.

## Supplemental information

**Supplemental Table 1 Primers used in this study**

**Supplemental Table 2 Off target analyses of base editors**

**Supporting information 1 Sequence of A3A-RAD51DBD**

## Notes

### Competing Interest Statement

The authors have declared no competing interest.

