## Supplementary Information for "Base editors for citrus gene editing"

**Supplemental Table 1 Primers used in this study**

| Oligo name | Sequence (5' to 3') | |
|---|---|---|
| ABE8-F1 | GAGAACACGGGGGACTCTAGAATGAAACGGACAGCCGACGGAAG | For ABE construct |
| ABE8-R1 | AGTTAGTTCCGATAGCCAGGCCGATGCTGTACTTCTT | For ABE construct |
| ABE8-F2 | GGCCTGGCTATCGGAACTAACTCTGTGGGATGG | For ABE construct |
| ABE8-R2 | TCTTCTTGGGCTCGAATTCATCAGCCCTTGAATCACCACCGAG | For ABE construct |
| CmY-F2 | TAAAACAATATTATCCCTGCAGGCGATCGGCGCGCCAGATTTGC | For CmYLCV promoter |
| CmY-R2 | ctgtccgtttcatTCTAGAAAGCTTAGCTCTTACCTGTTTTCGT | For CmYLCV promoter |
| CmY-R3 | AAGCTTAGCTCTTACCTGTTTTCGTC | For CmYLCV promoter |
| LOBBE-F1 | tgcaGTTTATATAGAGAAAGGAAA | For TATA box gRNA |
| LOBBE-R1 | aaacTTTCCTTTCTCTATATAAAC | For TATA box gRNA |
| UGI-F1 | TTCAAGGGCTGATgaattcAAGAGACCCGCAGCAACCAAGAA | For CBE UGI |
| UGI-R1 | tcttcttgggctcgaattGGCTCCCCCCCGACAGCATCTT | For CBE UGI |
| ALS-F | ATGGTCTCGTGCACAGGTCCCTCGGAGGATGATGTTTAAGAGCTATGCT GGAAACA | For ALS gRNAs |
| ALS-R | TaGGTCTCCAAACATCATCCTCCGCGGGACCTGTGCACCAGCCGGGAATC GAAC | For ALS gRNAs |
| LOBpro-F1 | CGATAAAATTCACCTCCATGTAATT | For TATA box genotyping |
| LOBpro-R1 | GAAGCGTGGAGAAGATTGAGAAG | For TATA box genotyping |
| CsALSgt-F1 | ACGAACAAGGCGGCATCTTCGC | For ALS genotyping |
| CsALSgt-R1 | CAATTTAATCGGCTGGTTCCAATTGG | For ALS genotyping |
| CsALSgt-F2 | CGCCACCAACCTGGTCAGCG | For ALS genotyping |
| CsALSgt-R2 | CCTGGCCGCCCAGATGTTGC | For ALS genotyping |
| Cas9gt-F1 | AACTAACTCTGTGGGATGGGCTGT | For Cas9 PCR |
| Cas9gt-R1 | TCTCGTGGTATGCCACCTCATCAA | For Cas9 PCR |
| Cs3g05060-F1 | AGATCAACATCTCAAAGCAGAGCA | For ABE off-target |
| Cs3g05060-R1 | GCCTATTCTTCTGAGTTCCTTGG | For ABE off-target |

| | | |
|---|---|---|
| Cs9g16440-F1 | ATGAGATCAACATCTCAAAGCAG | For ABE off-target |
| Cs9g16440-R1 | TTCCTCTTTGATGCTCACCTTC | For ABE off-target |
| Offtarget-F1 | CTTTTCCTAATATAGCCAGATTCAC | For ABE off-target |
| Offtarget-R1 | ATACATAAATCCGGATACGAAGTC | For ABE off-target |
| Offtarget-F2 | CCCTCTGCAAGGACACTGTCC | For ABE off-target |
| Offtarget-R2 | AGAAGTCATACCTATACCCGATC | For ABE off-target |
| Offtarget-F7 | AAAAAACAGAACTTGAGAAAGCGAA | For ABE off-target |
| Offtarget-R7 | GAATCTTGAATCTCTTGGATCATTA | For ABE off-target |
| Cs4g19110-F1 | GGTGGTCATTAAGGCCTGCAG | For ABE off-target |
| Cs4g19110-R1 | CATGACAACATAAAACCTGTTGAA | For ABE off-target |
| Cs2g08250-F1 | TCAACACACACAAGGAGAAGAGA | For ABE off-target |
| Cs2g08250-R1 | GCAACAGGTTGAGACTTTTGAAT | For ABE off-target |
| Cs5g30880-F1 | ATTCAATCTGTTGATCAATGTGATAA | For ABE off-target |
| Cs5g30880-R1 | AGCTTGTGGATAAAAGATAACAAGA | For ABE off-target |
| Cs6g18850-F1 | TTAATTAAACTTATTTAGTGGATCT | For ABE off-target |
| Cs6g18850-R1 | GGCAATTCAGCAACATCCTGC | For ABE off-target |
| Offtarget3-F1 | TAGAAGGATATAACGGTAATTACAC | For ABE off-target |
| Offtarget3-R1 | GTTTAAATAGCTTAACTCTTCATGG | For ABE off-target |
| Offtarget6-F1 | ACAACTGATTCTTTATTCTAGTAGG | For ABE off-target |
| Offtarget6-R1 | TCCCTATACTTATACATGGCAGC | For ABE off-target |
| Offtarget11-F1 | CCCTCCCTTAATTTCATTTCATAC | For ABE off-target |
| Offtarget11-R1 | ACCATATCACATGTCACTCCATC | For ABE off-target |
| Offtarget13-F1 | TGCTGGTTTGTCTGTTGCGCAA | For ABE off-target |
| Offtarget13-R1 | CAGCTTTTGTCACGTAACCTCC | For ABE off-target |
| CsALSoff-F | GGCCCAAAATCCAAAACAAAGACC | For CBE off-target |
| CsALSoff-R | GAAAAGAAGAGGGTTTTGTCAATCC | For CBE off-target |

### Supplemental Table 2 Off target analyses of base editors

| | Sequence | Locus | Gene | Region | Off-target edits in Hamlin mutant | Off-target edits in Grapefruit mutant |
|---|---|---|---|---|---|---|
| # | **For LOB1:** | | | | | |
| 1 | ATTTATATAAAGAAAAAAAAATGG | chrUn:+30450255 | | Intergenic | No | No |
| 2 | GTTTATAAAAAGAAAAAAAAAAGG | chr1:-8435697 | | Intergenic | No | No |
| 3 | GTTTAAAAAAAAAAAGGAAAAGG | chr1:-26798360 | | Intergenic | No | No |
| 4 | TTATATATATAGAAAGAAAAAGG | chr2:+4106970 | | Intergenic | No | No |
| 5 | TTATATGTAGAAAAAGGAAAAGG | chr3:+6405972 | Cs3g05060 | intron | No | No |
| 6 | TTATATGTAGAAAAAGGAAAAGG | chr9:-15859965 | Cs9g16440 | intron | No | No |
| 7 | GATTATATATAGAAGAGAAATGG | chr2:-30108859 | | Intergenic | No | N.A. |
| 8 | GTTTATAGAAGAAAAGGAAAAGG | chr6:+18869249 | Cs6g18850 | intron | No | No |
| 9 | GTTTGTACATGGAAAGGAAAAGG | chr7:-7084189 | | Intergenic | No | No |
| 10 | CTTTAAATTAAGAAAGGAAAAGG | chr5:-32692617 | Cs5g30880 | intron | No | No |
| 11 | GTATATGGAGAGAAAGAAAAAGG | chr2:-4970243 | Cs2g08250 | utr | No | No |
| 12 | ACTTATATGGAGGAAGGAAATGG | chr4:-18596584 | Cs4g19110 | CDS | No | No |
| | For CsALS: | | | | | |
| 1 | TTGGTGCCTCGGAGGGTGATGGG | chr9:-1639636 | | Intergenic | No | No |

Note: N.A., not amplified

**Supporting information 1 Sequence of A3A-RAD51DBD**

A3A-RAD51DBD

SV40 NLS

A3A

Linker

RAD51-DBD

```
CCCAAGAAGAAGCGTAAAGTTGAGGCAAGCCCAGCTTCAGGCCCCAGACACCTCATGGACCCACACATCT
TCACATCAAATTTCAATAATGGAATTGGGAGACATAAAACATATCTATGCTACGAAGTGGAGCGTCTCGA
TAATGGCACCAGCGTCAAGATGGACCAACATCGTGGATTTCTGCATAATCAAGCCAAAAACCTGCTATGT
GGGTTTTACGGACGACATGCAGAGTTACGTTTTCTTGACCTCGTGCCTTCACTACAGCTTGATCCCGCCC
AAATATATCGAGTTACCTGGTTTATTTCCTGGTCTCCATGTTTCTCTTGGGGGTGTGCTGGGGAAGTCCG
AGCCTTTTTACAGGAGAATACACATGTCAGGTTGAGGATCTTTGCTGCCAGAATCTATGATTATGATCCC
CTTTACAAGGAAGCACTCCAGATGTTAAGAGACGCTGGCGCACAAGTGAGCATTATGACTTATGACGAGT
TCAAACATTGCTGGGATACCTTTGTCGATCATCAAGGATGTCCTTTTCAGCCTTGGGACGGTCTGGACGA
ACATTCCCAGGCATTAAGCGGACGTTTGCGTGCTATACTTCAAAATCAAGGTAACTCCGGCGGGTCATCA
GGTGGCAGCAGTGGATCTGAGACCCCCGGTACTTCTGAATCAGCAACTCCTGAATCCTCTGGTGGCTCTT
CTGGGGGCAGCGCAATGCAAATGCAGCTTGAAGCAAACGCAGATACTTCCGTTGAAGAAGAGTCTTTCGG
ACCCCAGCCAATATCCCGACTTGAGCAGTGCGGCATAAATGCTAATGACGTAAAAAAACTGGAGGAGGCT
GGGTTCCATACTGTCGAAGCAGTGGCCTACGCACCTAAGAAAGAATTAATAAATATCAAGGGCATATCAG
AGGCAAAGGCTGACAAAATACTGGCTGAGGCCGCCAAGTTAGTCCCTATGGGATTTACCACTGCCACCGA
GTTCCATCAGCGTCGATCAGAGATTATACAAATCACAACTGGATCCAAAGAACTAGATAAACTGTTACAA
gtcgactccggaggatctag
```
